# Genomic diversity and hotspot mutations in 30,983 SARS-CoV-2 genomes: moving toward a universal vaccine for the “confined virus”?

**DOI:** 10.1101/2020.06.20.163188

**Authors:** Tarek Alouane, Meriem Laamarti, Abdelomunim Essabbar, Mohammed Hakmi, EL Mehdi Bouricha, M.W. Chemao-Elfihri, Souad Kartti, Nasma Boumajdi, Houda Bendani, Rokia Laamarti, Fatima Ghrifi, Loubna Allam, Tarik Aanniz, Mouna Ouadghiri, Naima El Hafidi, Rachid EL Jaoudi, Houda Benrahma, Jalil Elattar, Rachid Mentag, Laila Sbabou, Chakib Nejjari, Saaid Amzazi, Lahcen Belyamani, Azeddine Ibrahimi

## Abstract

The COVID-19 pandemic has been ongoing since its onset in late November 2019 in Wuhan, China. Understanding and monitoring the genetic evolution of the virus, its geographical characteristics, and its stability are particularly important for controlling the spread of the disease and especially for the development of a universal vaccine covering all circulating strains. From this perspective, we analyzed 30,983 complete SARS-CoV-2 genomes from 79 countries located in the six continents and collected from December 24, 2019, to May 13, 2020, according to the GISAID database. Our analysis revealed the presence of 3,206 variant sites, with a uniform distribution of mutation types in different geographic areas. Remarkably, a low frequency of recurrent mutations has been observed; only 169 mutations (5.27%) had a prevalence greater than 1% of genomes. Nevertheless, fourteen non-synonymous hotspot mutations (> 10%) have been identified at different locations along the viral genome; eight in ORF1ab polyprotein (in nsp2, nsp3, transmembrane domain, RdRp, helicase, exonuclease, and endoribonuclease), three in nucleocapsid protein and one in each of three proteins: spike, ORF3a, and ORF8. Moreover, 36 non-synonymous mutations were identified in the RBD of the spike protein with a low prevalence (<1%) across all genomes, of which only four could potentially enhance the binding of the SARS-CoV-2 spike protein to the human ACE2 receptor. These results along with mutational frequency dissimilarity and intra-genomic divergence of SARS-CoV-2 could indicate that the SARS-CoV-2 is not yet adapted to its host. Unlike the influenza virus or HIV viruses, the low mutation rate of SARS-CoV-2 makes the development of an effective global vaccine very likely.

## Introduction

The year 2019 ended with the appearance of groups of patients with pneumonia of unknown cause. Initial evidence suggested that the outbreak was associated with a seafood market in Wuhan, China, as reported by local health authorities [1]. The results of the investigations led to the identification of a new coronavirus in affected patients [2]. Following its identification on the 7th of January 2020 by the Chinese Center for Disease Control and prevention (CCDC), the new virus and the disease were officially named SARS-CoV-2 (for Severe Acute Respiratory Syndrome CoronaVirus-2) and COVID-19 (for Coronavirus Disease 19), respectively, by the World Health Organization (WHO) [3]. On March 11, 2020, WHO publicly announced the SARS-CoV-2 epidemic as a global pandemic.

This virus is likely to remain and continue to spread unless an effective vaccine is developed or a high percentage of the population is infected in order to achieve collective immunity. The development of a vaccine is a long process and is not guaranteed for all infectious diseases. Indeed, some viruses such as influenza and HIV have a high rate of genetic mutations, which makes them prone to antigenic leakage [4,5]. It is therefore important to assess the genetic evolution of the virus and more specifically the regions responsible for its interaction and replication within the host cell. Thus, identifying the conserved and variable regions of the virus could help guide the design and development of anti-SARS-CoV-2 vaccines.

The SARS-CoV-2 is a single-stranded positive-sense RNA virus belonging to the genus *Betacoronavirus*. The genome size of the SARS-CoV-2 is approximately 30 kb and its genomic structure has followed the characteristics of known genes of the coronavirus [6]. The ORF1ab polyprotein is covering two-thirds of the viral genome and cleaved into many nonstructural proteins (nsp1 to nsp16). The third part of the SARS-CoV-2 genome codes for the main structural proteins; spike (S), envelope (E), nucleocapsid (N) and membrane (M). In addition, six ORFs, namely ORF3a, ORF6, ORF7a, ORF7b, ORF8, and ORF10, are predicted as hypothetical proteins with no known function [7].

Protein S is the basis of most candidate vaccines; it binds to membrane receptors in host cells via its RBD and ensures a viral fusion with the host cells [8]. Its main receptor is the angiotensin-converting enzyme 2 (ACE2), although another route via CD147 has also been described [9,10]. The glycans attached to S protein assist the initial attachment of the virus to the host cells and act as a coat that helps the virus to evade the host’s immune system. In fact, a previous study has shown that glycans cover about 40% of the surface of the spike protein. However, the ACE2-RBD was found to be the largest and most accessible epitope [11]. Thus, it may be possible to develop a vaccine that targets the spike RBD provided it remains accessible and stable over time. Hence the importance of monitoring the introduction of any mutation that could compromise the potential effectiveness of a candidate vaccine.

This study aims to deepen our understanding of the intra-genomic diversity of SARS-CoV-2, by analyzing the mutational frequency and divergence rate in 30,983 genomes from six geographic areas (Africa, Asia, Europe, North and South America, and Oceania), collected during the first five months after the onset of the virus. These analysis generate new datasets providing a repository of genetic variants from different geographic areas, with particular emphasis on recurrent mutations and their distribution along the viral genome as well as estimating the rate of intraspecific divergence while evaluating the adaptation of SARS-CoV-2 to its host and the possibility of developing a universal vaccine.

## Results

### Diversity of genetic variants of SARS-CoV-2 in six geographic areas

A total of 30,983 SARS-CoV2 genomes from 79 countries in six geographic areas (Africa, Asia, Europe, North and South America, and Oceania) included in this analysis. According to the GISAID database, the date of collection of the strains was within the first five months following the onset of SARS-CoV-2. A total of 3,206 variant sites were detected compared to the reference genome Wuhan-Hu-1/2019. Then, we analyzed the type of each mutation, highlighting the prevalence of these mutations both in all genomes (worldwide) and in each of the geographic areas studied (**Figure 1**). Worldwide, 67.96% of mutations had a non-synonymous effect (64.16% have missense effects, 3.77% produce a gain or loss of stop codon and 0.33% produce a loss of start codon), 28.60% were synonymous, while 3.43% of the mutations were localized in the intergenic regions, mainly in the untranslated regions (UTR). Likewise, the comparison between the six geographic areas shows a similar trend with a uniform distribution of the prevalence of each type of mutation.

**Figure 1:**
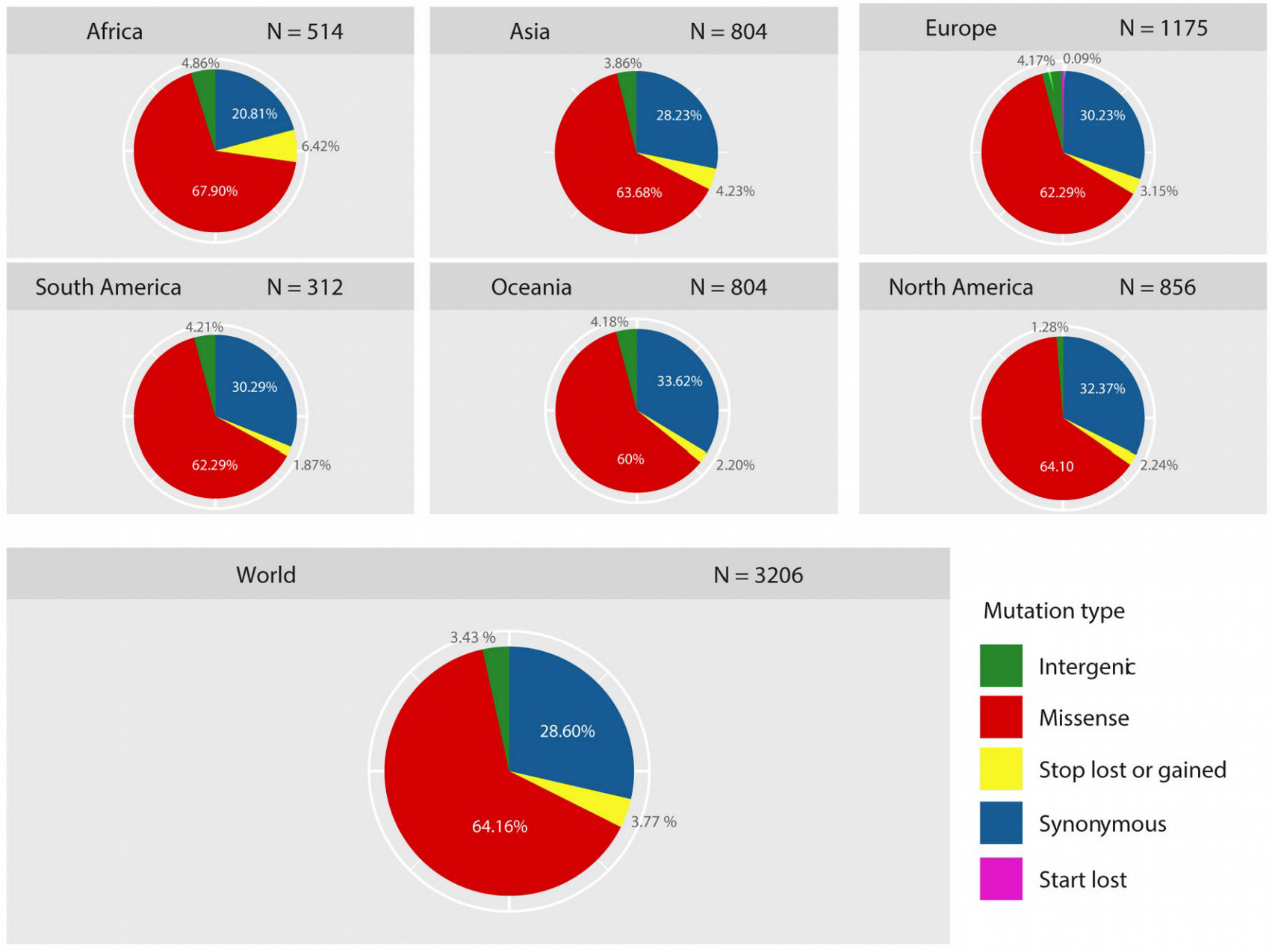
Prevalence and distribution of types of mutations in different geographic regions. Pie charts showing the global and continent-stratified distribution of the mutation types identified in the 30,983 SARS-CoV-2 genomes. The prevalence of each type of mutation is uniform across the six geographic areas and missense mutations were the most frequent type. Color codes represent the type of mutations.

The frequency of mutations in the six geographic areas was estimated by normalizing the number of genomes carrying a given mutation per the total number of genomes recovered by geographic area. Only 169 (5.27%) variant sites were found with a frequency greater than 0.01 (**Figure 2A**), and were distributed in six geographic areas as follow: 69 in Oceania, 65 in Africa, 54 in Asia, 31 in Europe, 43 in North America, and 43 in South America. Focusing on non-synonymous mutations (with a frequency> 0.01), 3.34% (n = 107) of the total mutations were identified (**Figure 2B**).

**Figure 2:**
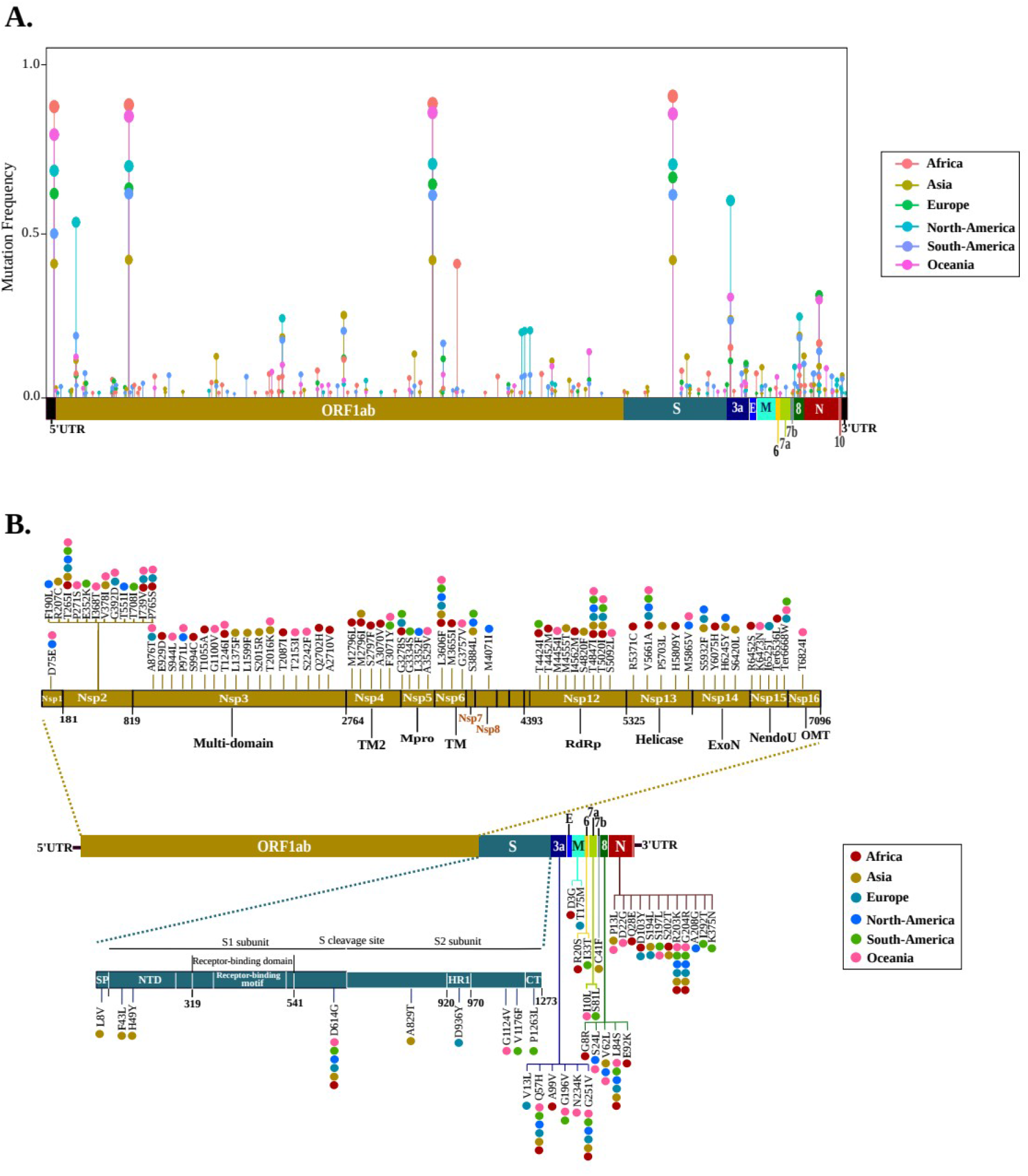
Distribution of recurrent mutations across the SARS-CoV-2 genome. **(A)** Lollipop plot illustrating the location of mutations with a frequency greater than 0.01 of the total genomes of each geographic area. All types of mutations are included (non-synonymous, synonymous and intergenic). The mutation frequency was estimated for each of them, by normalizing the number of genomes harbored a given mutation in a geographic area, per the total number of genomes recovered by geographic area. **(B)** Schematic representation illustrating the distribution of nonsynonymous mutations (with a frequency > 0.01) along the viral genome. Amino acid mutations are shown by vertical lines. Colored dots represent geographic areas.

The polyprotein ORF1ab contained approximately two thirds of these muations (63.55%; n = 68) and distributed in thirteen non-structural proteins; nsp3-Multi-domain: 15.89%, nsp2: 11.21%, nsp12-RNA-dependent RNA polymerase (RdRp): 8.41%, nsp4-transmembrane domain-2 (TM-2): 4.67%, nsp13-helicase: 4.67%, nsp15-endoribonuclease (NendoU): 4.67%, nsp5-main proteinase (Mpro): 3.74%, nsp14-exonuclease (ExoN): 3.74%, nsp6-TM: 2.80%, nsp1: 0.93%, nsp7: 0.93%, nsp8: 0.93% and nsp16-2’-O-ribose methyltransferase (OMT): 0.93%. The rest (36.45%) were distributed in eight proteins, including N (11.21%), S (8.41%), ORF3a (5.61%), ORF8 (4.67%), M (1.87%), ORF6 (1.87%), ORF7a (1.87%) and ORF7b (0.93%).

Comparative analysis of these non-synonymous mutations shows only nine that have been shared in the six geographic areas,: T265I (nsp2), L3606F (in nsp6-TM) T4847I (in nsp12-RdRp), D614G (in S), R203K-G204R (in N), Q57H-G251V (in ORF3a) and L84S (in ORF8). It is also interesting to note that none of the nine non-synonymous mutations (> 0.01) of S protein was localized in RBD. The 36 non-synonymous mutations (35 with a missense effect and 1 with a stop gain effect) found in this area had a low frequency (< 0.01) across all genomes. Among them, only two mutations were shared between genomes of different geographic areas; the V367F mutation was identified in in Europe, Asia, and North America, the V367F mutation has been identified in Europe, Asia and North America, while P491L in Asia and Oceania.

### Geographical distribution of the SARS-CoV-2 hotspot mutations

Comparative genomic analysis of each geographic area revealed fourteen non-synonymous mutations with a frequency greater than 0.1 and considered as hotspot mutations (**Figure 3**). Eight mutations of them were found in the ORF1ab polyprotein, distributed in seven regions coding for nsp2 (T265I), nsp3-Multi-domain (T2016K), nsp6-TM (L3606F), nsp12-RdRp (T5020I and T4847I), nsp13-helicase (M5865V), nsp14-ExoN (D5932T) and nsp15-NendoU (Ter6668W). Moreover, three mutations in N protein (R203K, G204R, and P13L) and one in each of the three proteins; S (D614G), ORF3a (Q57H) and ORF8 (L84S).

**Figure 3:**
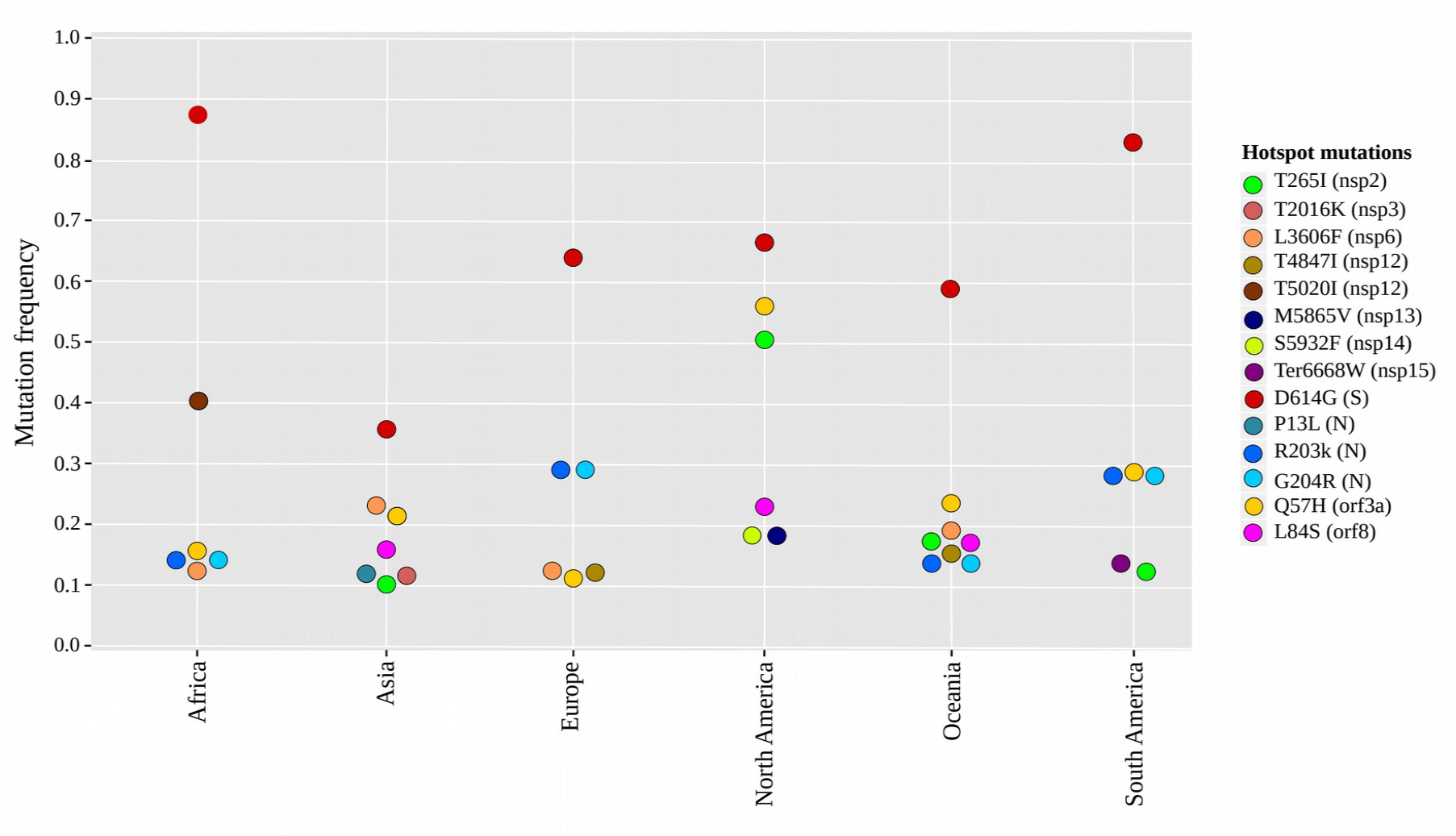
Frequencies of recurrent hotspot mutations per geographic area. Distribution of fourteen non-synonymous mutations with a frequency> 0.1 of the genomes subdivided into six geographical areas; Africa (n = 6), Asia (n = 7), Europe (n = 6), North America (n = 6), Oceania (n = 8), South America (n = 6). The locations of mutations in viral proteins with their color codes are indicated in the legend.

Different patterns of these non-synonymous hot spot mutations were observed between the six geographic regions. Only two mutations were common in the six geographical regions: the high-frequency mutation D614G (in S) and the Q57H mutation (in ORF3a). Seven mutations were more frequent in a single geographic region, including two mutations T2016K (in nsp3-Multi-domain) and P13L (in N) in Asia, two mutations M5865V (in nsp13-helicase) and D5932T (in nsp14-ExoN) in North America, one T5020I (in nsp12-RdRp) in Africa, one T4847I (in nsp12-RdRp) in Europe and one Ter6668W (in nsp15-NendoU) in South America. However, the other five non-synonymous hotspot mutations were variable between the six geographical regions, including two R203K and G204R (in N) that were particularly predominant in Africa, Europe, South America, and Oceania. Whereas, L3606F (in nsp6-TM) was common in Africa, Asia, Europe, and Oceania. Thus, L84S (in ORF8) was found in Asia, North America, and Oceania. In addition, T265I (in nsp2) was frequent in Asia, North America, South America, and Oceania.

### The distribution of hotspot mutation patterns of SARS-CoV-2 over time

In order to systematically investigate the appearance of each non-synonymous hotspot mutations, we analyzed each genome in each geographical area over time, classifying them per date of sample collection (according to GISAID database). We have noticed that the distribution of the fourteen hotspot mutations patterns is dynamic during the propagation of SARS-CoV-2 in different geographic regions.

Among the fourteen hotspot mutations, four first appeared in January 2020, seven in-February 2020, and three in the first two weeks of March 2020, (**Figure 4**).

**Figure 4:**
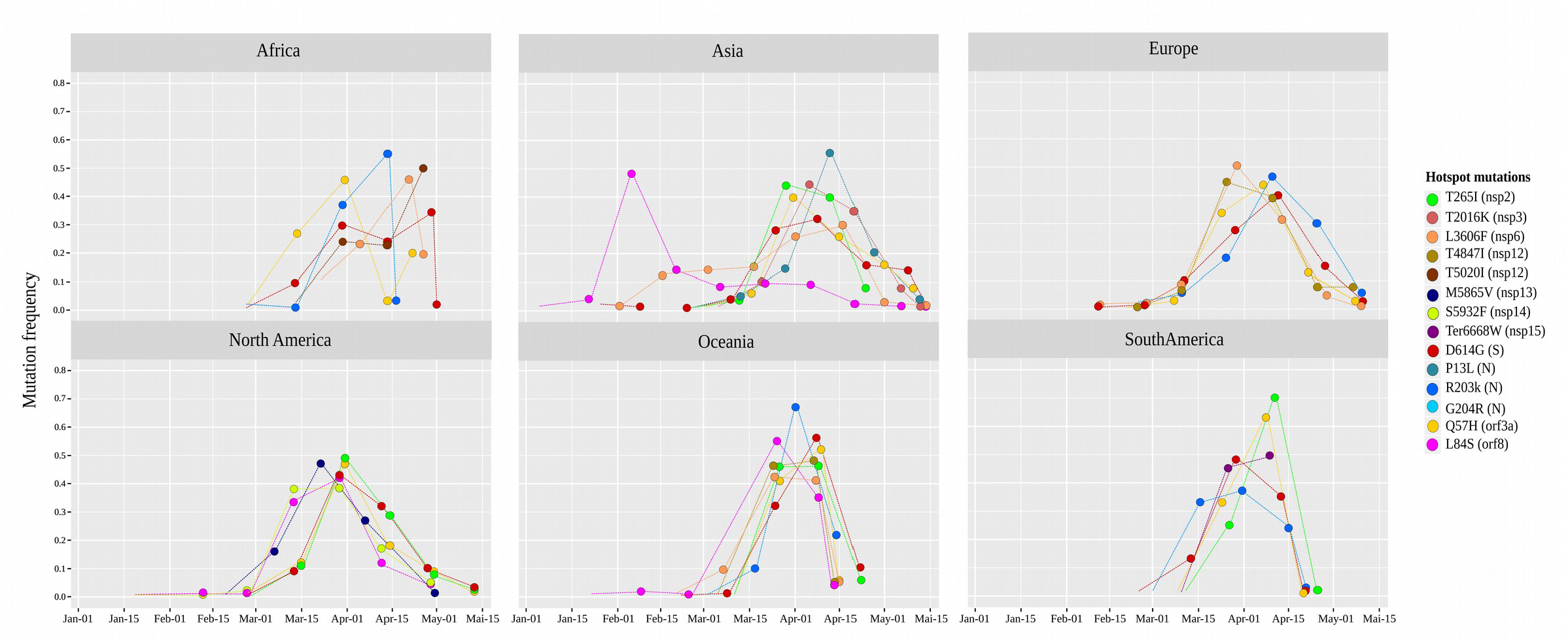
Tracking hotspot mutations over time per geographic area. Hotspot mutation frequencies was plotted for each of them over a period of 15 days from their appearance in each geographic area, first by normalizing the number of genomes harboring a given mutation in a period of 15 days, by the total number genomes harboring this mutation. The X axis represents the time measured in 15 days and the Y axis represents the frequencies of the genomes harboring the hotspot mutations.

The high-frequency mutation D614G (in S), appeared on January 24th, 2020, in Asia (China), next it was observed in Europe (Germany) after four days, and then it gained predominance over time due to its spread to many European countries. In late February, this mutation was also reported in Africa (Nigeria and Senegal), South America (Brazil), North America (Mexico and USA), and Oceania (Australia).

The L84S mutation (in ORF8) appeared on January 05th, 2020 in Asia (China), then the emergence of this mutation was observed in the second half of February in the North American and Oceanic regions, particularly in the USA and Australia, respectively. For L3606F (in nsp6-TM) emerged in the third week of January in Asia (China), then propagated in late January in Europe (France and Italy), then in the third week of February and March 2020, in Oceania (Australia) and Africa (DRC and Senegal), respectively. While the mutation D5932T (in nsp14-ExoN) appeared on January 19th, 2020, particularly in North America (USA).

Among the hotspot mutations that first appeared in February 2020, three within the N protein gene (R203K, G204R, and P13L) gained their predominance during the COVID-19 pandemic. Indeed, the double mutations R203K-G204R emerged on the same date in different geographic areas; February 24th in Europe, February 27th in Africa (in Nigeria), March 2nd in Oceania (New Zealand) and March 04th, 2020, in South America (Brazil). Meanwhile the P13L appeared in late February, particularly in Asia (Korea). We also noticed an increase in the frequency of the Q57H mutation (in ORF3a) in late February in Africa (Senegal), Europe (France and Belgium) and North America (USA and Canada). It was also observed in the three other regions at the beginning of March in Asia (Taiwan), Oceania (Australia) and South America (Brazil). Thus, we found that the T265I (in nsp2) mutation first appeared in late February in both Asia (Taiwan) and North America (USA), while it was spread at the beginning of March in South America (Brazil) and Oceania (Australia). In addition, T4847I (in nsp12-RdRp) gained predominance from the second week of February in Europe (United Kingdom), then after one month spread in Oceania (Australia). While the frequency of the M5865V mutation (in nsp13-helicase) has increased particularly in North America (USA).

For the three hotspot mutations that appeared during the first two weeks of March 2020, were found with a frequency greater than 0.1 in genomes belonging to a single geographic region, including T2016K (in nsp3-Multi-domain), Ter6668W (in nsp15-NendoU) and T5020I (in nsp12-RdRp) in Asia (Taiwan), South America (Chile) and Africa (Senegal), respectively.

### Mutagenesis of D614G and impact of RBD mutations on the binding ability of spike to ACE2

As shown the **Figure 5**, the non-synonymous D614G mutation did not have an impact on the two- or three-dimensional structure of the spike glycoprotein. However, D614 residue in the wild type spike is involved in three hydrogen bonds; one with A647 in the same subunit (S1) and two bonds with THR-859 and LYS-854 located at S2 subunit of the adjacent protomer (**Figure 5A**). The substitution of D614 by G in the mutant spike resulted in the loss of the two hydrogen bonds with THR-859 and LYS-854 in the S2 subunit of the adjacent protomer (**Figure 5B**). Such modification could result in a weak interaction between S1 and S2 subunits and thus increases the rate of S1/S2 cleavage, which will improve the virus entry to host cells.

**Figure 5:**
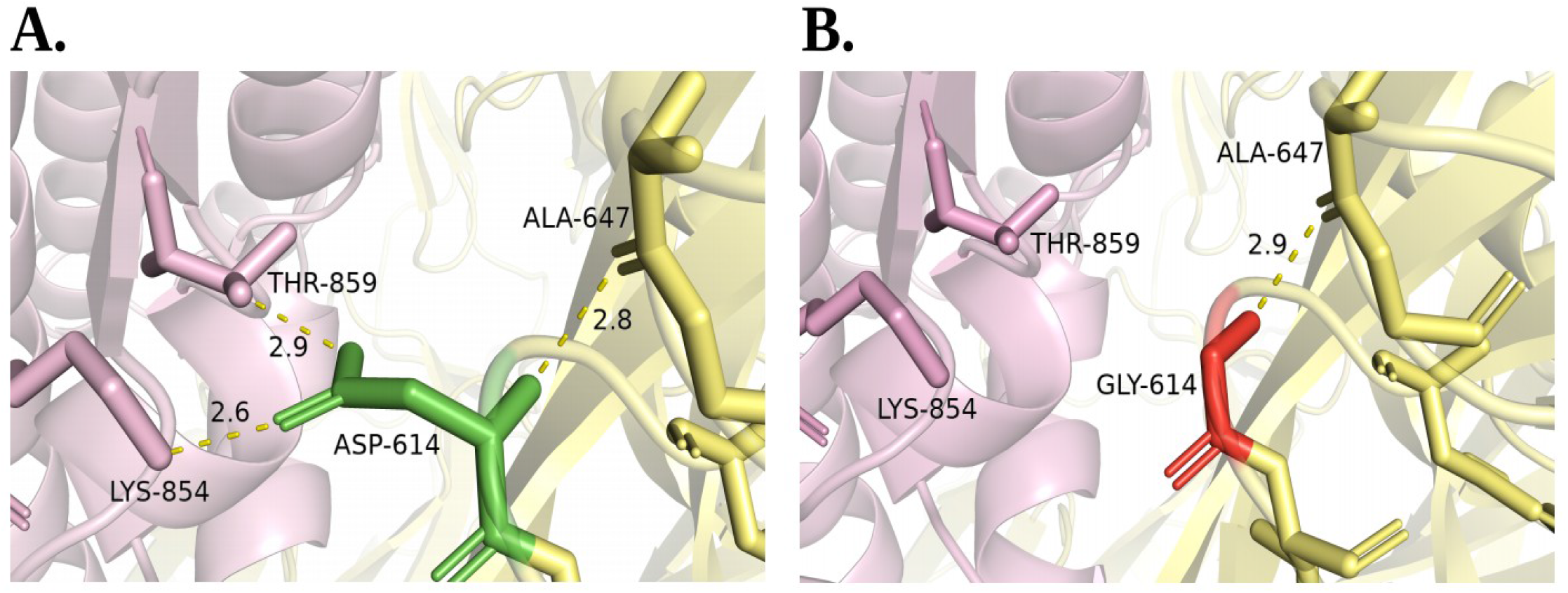
Comparison of spike wild type residue ASP-614 **(A)** and the mutated GLY-614 **(B)**. ASP-614 (green sticks) in subunit S1 (yellow) is involved in two hydrogen bonds with THR-859 and LYS-854 from the S2 subunit (pink). The substitution of ASP by GLY at position 614 causes the loss of the two hydrogen bonds between S1 subunit and THR-859 and LYS-854 in the S2 subunit (pink).

To evaluate the effect of RBD mutations on the binding affinity of the spike protein to ACE2, the MM-GBSA method was employed to calculate the binding affinity of 35 spike mutants to ACE2 (except for the stop-gain mutation). Four mutations potentially enhance the binding affinity of spike/ACE2 complex by more than −1.0 kcal/mol, while nine were shown to potentially reduce its affinity by more than 1.0 kcal/mol (**Table 1**). However, the remaining 22 did not significantly affect the binding affinity of spike to ACE2.

**Table 1.** Impact of mutations on the binding affinity between spike protein RBD and ACE2, evaluated by MMGBSA binding free energy calculation (ΔG_Bind_).

| Mutations | $\Delta G_{\text{Bind}}$ (kcal/mol) | $\Delta \Delta G^1$ (kcal/mol) | Effect on Spike/ACE2 |
| --- | --- | --- | --- |
| V367F | -62.47 | 3.53 | Potentially decreased binding Affinity |
| S477N | -62.69 | 3.31 |  |
| R408I | -62.8 | 3.2 |  |
| V483A | -63.85 | 2.15 |  |
| A522S | -64.03 | 1.97 |  |
| G339D | -64.08 | 1.92 |  |
| N354D | -64.39 | 1.61 |  |
| K356N | -64.81 | 1.19 |  |
| H519Q | -64.84 | 1.16 |  |
| Wild Type | -66 | 0 | Wild type MMGBSA value |
| N440K | -67.88 | -1.88 | Potentially decreased binding Affinity |
| N450K | -67.88 | -1.88 |  |
| D364Y | -68.24 | -2.24 |  |
| S477R | -69.86 | -3.86 |  |

<sup>1</sup> binding free energy difference between mutated and wild type complexes.

### Clustering and divergence of SARS-CoV-2 genomes

This section aims to identify countries sharing similar trends in mutational frequencies. The clustering with Jaccard distance between the genomes of each country based on their mutational frequencies showed two main clusters (cluster 1 and cluster 2) (**Figure 6**), each subdivided into several sub-clusters (SCs) and these two clusters include the countries of the six continents. In addition, the countries in group 2 were closer to each other than those in cluster 1, demonstrating high genetic similarities between strains from these countries. We also observed fifteen SCs with a distance of less than 0.5 (**Fig 6, Table 2**), including ten SCs belonging to Cluster 2. Remarkably, most of the countries grouped in these fifteen SCs are geographically close, of which eleven SCs include countries of the same continent, especially Asia and Europe.

**Table 2.** Jaccard distance between countries based on their mutational frequencies. Only the distance less than 0.5 between countries is displayed.

| Cluster | Sub-Cluster | Countries | Jaccard distance | Geographic areas |
| --- | --- | --- | --- | --- |
| Cluster 1 | SC-1 | Brunei, Guam | 0.22 | Asia, Oceania |
| Cluster 2 | SC-2 | Kazakhstan, Georgia | 0.22 | Asia |
| Cluster 2 | SC-3 | Nigeria, Serbia, Croatia, Ireland, Peru | 0.26 | Africa, Europe, South America |
| Cluster 2 | SC-4 | Vietnam, Jordan | 0.27 | Asia |
| Cluster 2 | SC-5 | Sri Lanka, Kuwait | 0.30 | Asia |
| Cluster 2 | SC-6 | Greece, Portugal | 0.32 | Europe |
| Cluster 2 | SC-7 | Singapore, Thailand | 0.35 | Asia |
| Cluster 2 | SC-8 | Finland, Poland | 0.35 | Europe |
| Cluster 2 | SC-9 | Slovenia, Jamaica | 0.35 | Europe, North America |
| Cluster 2 | SC-10 | Denmark, Iceland | 0.36 | Europe |
| Cluster 2 | SC-11 | Germany, Russia | 0.36 | Europe |
| Cluster 1 | SC-12 | Hungary, Latvia | 0.41 | Europe |
| Cluster 1 | SC-13 | Chile, Brazil | 0.43 | South America |
| Cluster 1 | SC-14 | Iran, Pakistan | 0.43 | Asia |
| Cluster 2 | SC-15 | Netherland, Belgium, Austria | 0.49 | Europe |

**Figure 6:**
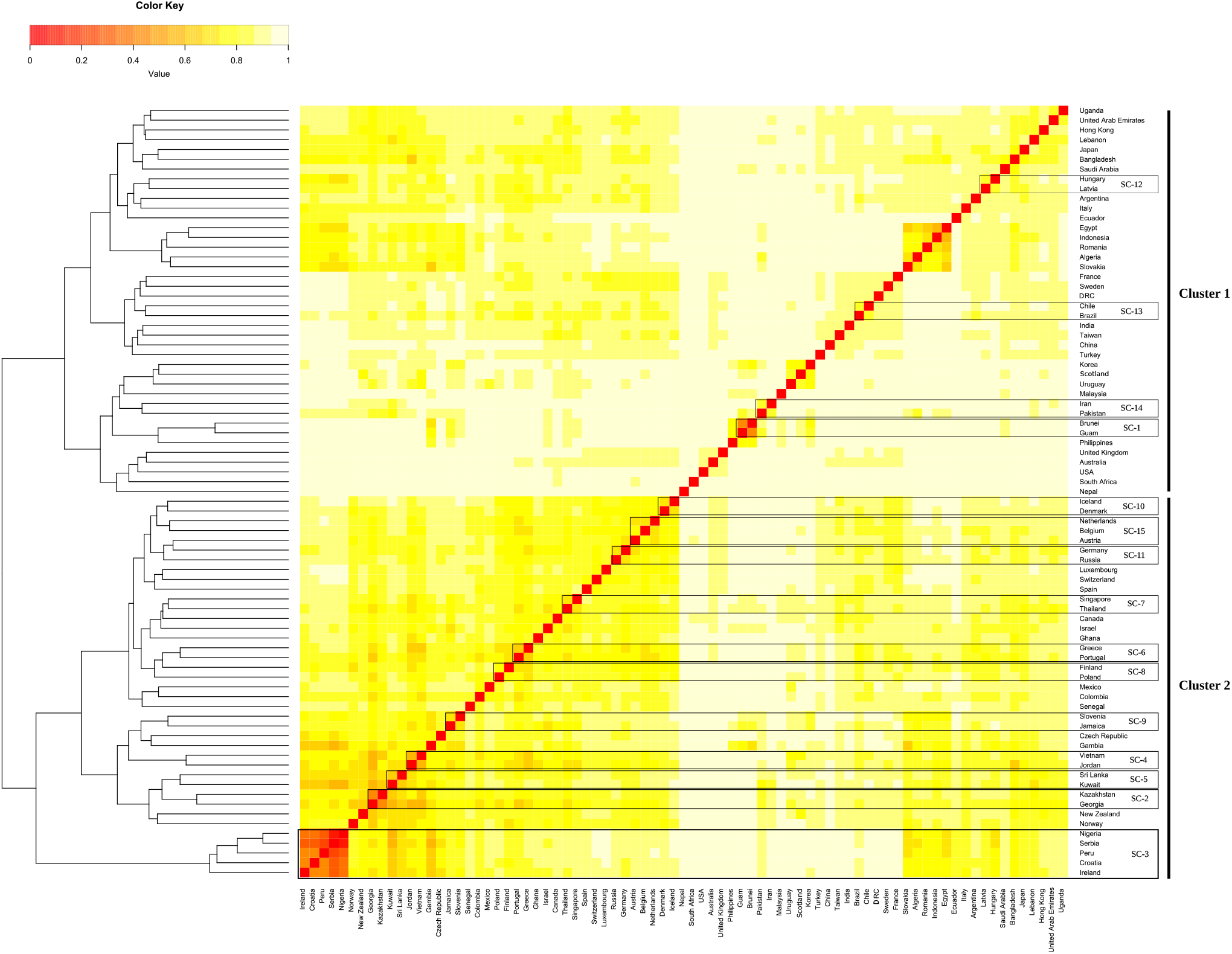
Dendrogram and Heatmap of the hierarchical analysis of clusters between countries. The distance matrix between these countries was calculated using Jaccard Distance, and the values ranged from 0 (light yellow) to 1 (red). Countries connected with a distance <0.5 were presented as a sub-clusters of SC-1 to SC-15.

Meanwhile, the intraspecific divergence of SARS-CoV-2 was also assessed in the genomes of each country compared to the genome reference Wuhan-Hu-1/2019. As shown in **Figure 7A** the overall circulating strains in more than 50 countries seem to have a divergence percentage of less than 0.1%, which indicates that the majority of SARS-CoV-2 genomes have developed less than 18 mutations in them.

**Figure 7:**
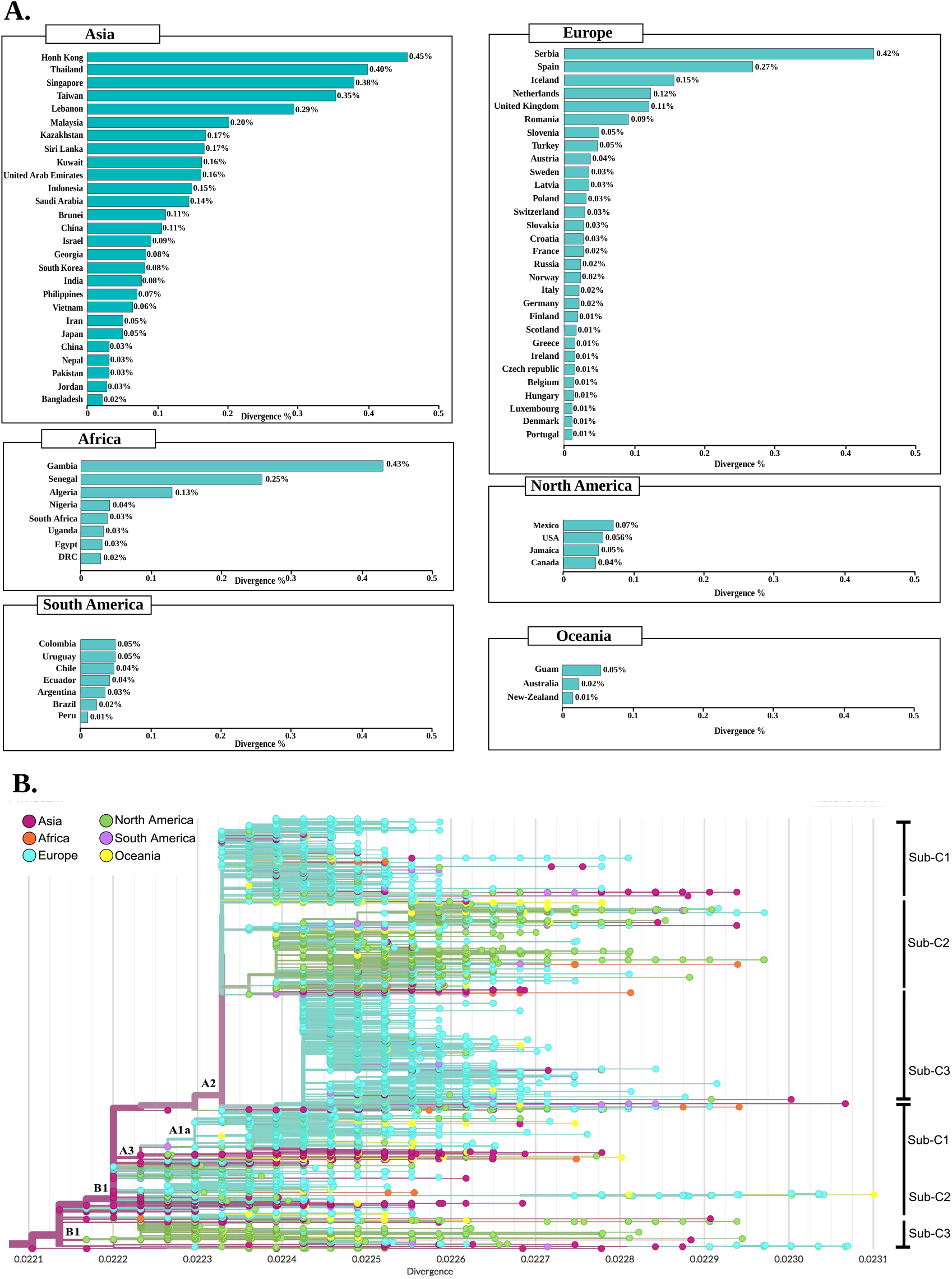
Divergence of SARS-CoV-2 genomes from different geographic areas compared with the genome reference Wuhan-Hu-1/2019. **(A)** The bar graph illustrating the divergence (measured in percentage) of the SARS-CoV-2 genomes of each country compared to the reference genome Wuhan-Hu-1/2019. The divergence calculation method is detailed in the Material and Method section. **(B)** The phylogenetic divergence tree of the 30,983 SARS-CoV-2 genomes grouped into six geographic regions. The length of the branches shows the divergence and the color codes indicate the six geographical areas.

The highest percentage of divergence in Asia, Europe, North America, South America, Oceania, and Africa was observed in Hong Kong (0.45%), Serbia (0.42%) Mexico (0.07%), Colombia (0.05%), Guam (0.05%) and Gambia (0.43%), respectively. While the lowest percentage was shown in Portugal (0.01), Canada (0.05%), Bangladesh (0.02%), Peru (0.01%), New Zealand (0.02%) and DRC (0.03%).

Moreover, the phylogenetic divergence tree (**Figure 7B**) shows that the highest rate was among genomes from Asia, followed by Europe, and North America. In Asia, most strains showed a divergence of 0.0221 to 0.0231. Likewise, European strains clustered between 0.0223 and 0.0231, while North American strains had a divergence of 0.0221 to 0.0230. Using the Nextstrain clade nomenclature, we can identify two main clades with different divergence profiles; first and most divergent “A2” clade, although the first strain observed was from China. This clade mainly contains genomes from Europe. The second “B1” clade appears to be less divergent and to a large extent includes Asian strains. Nevertheless, the genomes of Africa, North America, and South America have been scattered across the phylogenetic divergence tree without a specific coating.

The rate of divergence also varied within clades, A2 includes three subclades, the sub-c2 harboring the Q57H mutation, having a divergence of 0.0224 to 0.0229, and mainly includes strains from North America and North America. ‘Asia. The sub-c3 containing mostly European genomes sharing the R203K mutation, in this subclade a low rate of divergence was observed in all continents except Africa, while the greatest divergence was a strain from Taiwan (Asia) (> 0.0223).

On the other hand, clad 2 (B1) harbored mainly genomes from North America and Asia, while the high divergence in this clade observed in Europe (France) with 0.0231. The sub-c2 and sub-c3 of this clade appeared to be the most diverse with the lowest divergence in the United Kingdom and the highest in Australia.

### Phylogenetics and spatio-dynamics of SARS-CoV-2

The topology of the maximum likelihood phylogenetic tree (**Figure 8A**) shows a clear clustering: one cluster containing mainly Asian strains, while the second containing European strains with a specific clade sharing the D614G mutation. For each cluster, we identified different clades: cluster I containing two main clades A1a and B1 harboring mainly strains from Asia, North America, and Asia, Europe, respectively. However, cluster II harbored three clades: B2, A2, A2a without a specific pattern.

**Figure 8:**
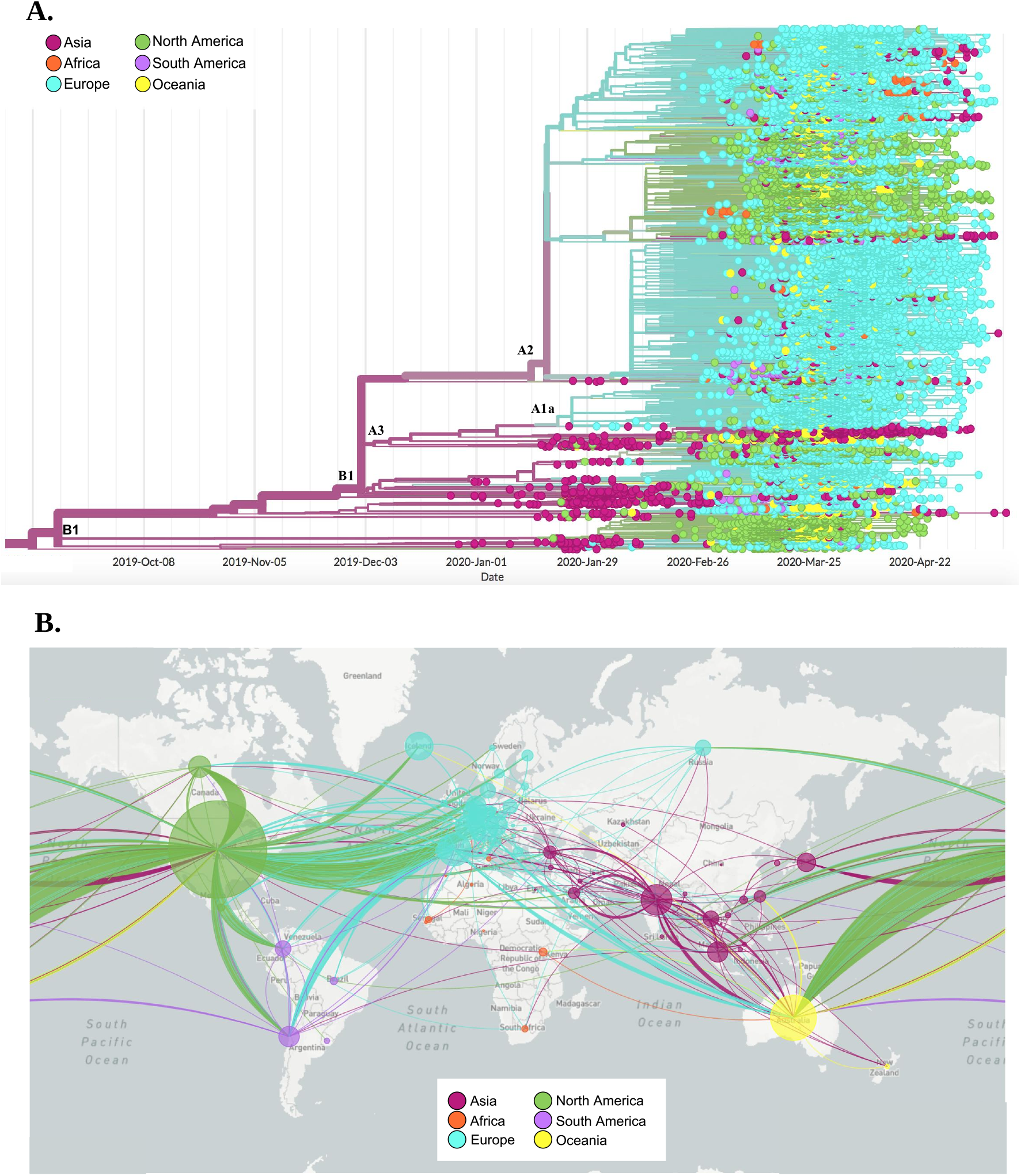
Phylogenetic tree and spatial dynamics of SARS-CoV-2. **(A)** Phylogenetic analysis of 30,983 SARS-CoV 2 genomes grouped into six geographic areas. The length of the branches represents the distance in time. **(B)** Phylodynamic analysis representing the propagation and evolution of 30,983 SARS-Cov-2 genomes in different geographic areas. The color codes represent the six geographic areas

The distribution of African genomes across the phylogenetic tree showed a close relationship with different continents. In the first clade, African genomes (mainly from South West Africa) clustered with Asia and showed the lowest divergence rate. Meanwhile, genomes clustering in the European clade sharing the three-pattern mutations mostly common in Europe: G28881A, G28882A, and G28883C.

The map (**Figure 8B**) shows the spatio-dynamics of SARS-CoV-2 and provides an insight into the viral strain’s potential geographical origin based on the sample used and displays a complex and interconnected network of strains. From these samples, strains from Asia appear to have diverged and resulted in other strains in all the investigated regions. European strains seem to have given rise to those in North America, South America and Africa, with multiple divergent strains within Europe itself. Similarly, with less frequency, strains from South and North America appear to be related to some divergent strains in Europe and Asia.

## Discussion

Due to the rapid spread and mortality rate of the new SARS-CoV-2 pandemic, the development of an effective vaccine against this virus is of a high priority [12]. The availability of the first viral sequence derived during the COVID-19 epidemic, Wuhan-Hu-1, was published on January 5th, 2020. From this date, numerous vaccination programs were launched [13,14]. Furthermore, drugs and vaccines should target relatively invariant and highly constrained regions of the SARS-CoV-2 genomes, in order to avoid drug resistance and vaccine escape [15]. For this, monitoring genomic changes in the virus are essential and play a pivotal role in all of the above efforts, due to the appearance of genetic variants, which could have an effect on the efficacy of vaccines. In this study, we investigated the genetic diversity of the 30,983 complete SARS-CoV-2 genomes isolated from six geographic areas.

Our results showed three different situations of the identified mutations; (i) the mutations that have developed and are gaining a predominance in the six geographic areas; (ii) mutations which were predominant only in certain geographic regions and (iii) mutations apparently expanding, but low in frequency in all isolates studied. From this third situation, it is interesting to note that a low rate of recurrent mutations was found across genomes, with only 5.27% of the total mutations having a frequency greater than 0.01. While 94.73% had a frequency of less than 0.01, of which 49.68% were single mutations (specific to a genome). These results indicate that, so far, SARS-CoV-2 has evolved through a non-deterministic process and that random genetic drift has played the dominant role in the spread of low-frequency mutations worldwide, suggesting that the SARS-CoV-2 is not yet adapted to its host, and epidemiological factors are particularly responsible for the global evolution of this virus. Although some hotspot mutations have increased their frequencies, they have systematically gained their predominance outside Asia. For example, the hotspot mutation Q57H (in ORF3a), on February 21st, 2020, has not yet been observed among isolates from China, while it has emerged in Europe and also spread in isolates from North America. Likewise, seven other mutation hotspots saw their frequencies in the SARS-CoV-2 genomes to be increased over time outside of Asia; including, double mutations R203K-G204R of protein N (in Europe, South America, and Oceania), M5865V of nsp13-helicase and D5932T of nsp14-ExoN (in North America), T5020I of nsp12-RdRp (in Africa), T4847I from nsp12-RdRp (in Europe and Oceania), Ter6668W from nsp15-NendoU (in South America).

It is interesting to note that the ORF10 protein was the only invariant protein in recurrent mutations. In our recent study [16], we reported the absence of this protein in certain SARS-CoV-2 genomes, which could explain a rate of low-frequency mutations of this protein. Without neglecting also its small size compared to other proteins.

A low rate of divergence was found in the genomes of each country compared to the reference genomes Wuhan-Hu-1/2019. Indeed, China was the source of the original variant, it had a high divergence rate among most of the countries in different geographies. This could suggest early and rapid transmission of infections to the worst-affected countries of Europe and America by variants genetically close to the original strain. The rapid transmission meaning a single source leading to multiple infections, thus giving the virus fewer life cycles to change. When the disease spreads slowly, the virus goes through several cycles of spread and is therefore likely to be more different from the original variant. This is consistent with a previous study describing a continued tendency for the virus to diverge over time [17]. In addition, accumulations of mutations have been shown to aid the adaptation of the virus to the human host by optimizing codon use through synonymous mutations [18]. Overall, this study showed an intraspecific SARS-CoV-2 divergence rate of less than 0.5% across the genomes of each country. Comparing all countries the high rate of discrepancy in Asian countries could be due to multiple sources of infection with different strains at the start of the epidemic. Likewise, intra-genomic clustering did not have a clear pattern regarding their geographic distributions, reflecting the effect of migration and globalization as previously indicated [19,20].

The phylogenetic tree confirms that disease transmission in Africa probably has three sources. The least divergent variants were grouped with Asia while the most divergent were grouped with Europe and North America. This distribution points to different sources of infection; countries in West and North Africa probably had their carriers from European countries. While South American genomes appear to originate from North America and Europe. While some strains from Oceania and Africa allow poor monitoring of the origin of the infection but show a close relationship with the genomes of Europe. Taken together, the North American and European genomes appear to be responsible for most of the spread of the disease.

As the virus spreads more widely around the world, it is important to follow the appearance of new mutations development that could compromise the effectiveness of a candidate vaccine. The SARS-CoV-2 S protein is the primary target of most vaccine candidates that are under development, due to its key role in mediating entry of the virus and its immunogenic trait [8,21]. Analysis of S protein in the SARS-CoV-2 genomes that we collected in this study revealed a hotspot mutation at position 614 that involves the change from a large amino acid residue (aspartic acid) to a small hydrophobic one (glycine). This mutation was most frequent in the genomes of six geographical areas. Several studies had already indicated its emergence [16,22–25]. The D614G mutation is proximal to the S1 cleavage domain of the spike glycoprotein [26]. Our finding showed that this mutation induces a loss of two hydrogen bonds between the S1 and S2 subunits of neighboring protomers and may, therefore, increase the cleavage rate of these subunits in the pre-fusion state of the spike protein to allow its conformational transition to the post-fusion state that is associated with membrane fusion during virus entry [27].

However, our structural modeling of this mutation has shown no substantial impact on the secondary or tertiary structure of the spike protein which makes the likelihood of this variant significantly affect a potential epitope in this region very low. From a clinical perspective, there is no evidence that this mutation affects the transmissibility and virulence of SARS-CoV-2 due to the lack of association between the D614G mutation and patient hospitalization status [28]. With this in mind, we believe that this will not present a serious issue for vaccine development and that a universal candidate vaccine for all circulating strains of SARS-CoV-2 may be possible.

On the other hand, the RBD of S protein, allows the virus to bind to the ACE2 receptor on host cells [29,30]. Mutations in this receptor are a likely route to escape recognition of antibodies, as described in other viruses [31,32]. In all of the genomes analyzed in this study, only 36 non-synonymous mutations in RBD have been identified. These mutations observed at low frequency (< 0.01) in all genomes, the calculated binding free energy of mutant RBDs of spike protein in complex with ACE −2 demonstrate that in 89% of the cases these mutations decrease or does not significantly affect the ability of the virus to bind to the host cells receptor. While only 11% potentially enhance the binding of the viral spike protein to ACE-2 and therefore could influence the pathogenicity of SARS-CoV-2.

SARS-CoV-2 has only been identified recently in the human population; a short time compared to the adaptive processes that could take years to occur. Although we cannot predict whether adaptive selection will be observed in SARS-CoV-2 in the future, the main conclusion is that the currently circulating SARS-CoV-2 viruses constitute a homogeneous viral population.

This study is a comprehensive report on the amino acid mutations that spread during the SARS-CoV-2 pandemic, from December 24th to May 13th. We can, therefore, be cautiously optimistic that, up to now, genetic diversity should not be an obstacle to the development of a broad protection SARS-CoV-2 vaccine. Therefore, ongoing monitoring of the genetic diversity of SARS-CoV-2 will be essential to better understand the host-pathogen interactions that can help inform drug and vaccine design.

## Materials and Methods

### Data collection

Full-length viral nucleotide sequences of 30,983 SARS-CoV-2 genomes were collected from the GISAID EpiCovTM (update: 26-05-2020) [33], belonging to the six geographical zones (according to GISAID database) and distributed in 79 countries as follow: 214 from Africa, 368 from South America, 1,590 from Oceania, 2,111 from Asia, 6,825 from North America and 19,875 from Europe. The genomes were obtained from samples collected from December 24, 2019, to May 13, 2020.

For each geographical area the collection date of the samples is from February 27^th^ to May 01st for Africa, December 24th to May 13th for Asia, January 23th to May 10^th^ for Europe, January 19th to May 12th for North America, February 25th to April 19th and January 24th to April 21st for Oceania.

### Variant calling analysis

Genome sequences were mapped to the reference sequence Wuhan-Hu-1/2019 (Genbank ID: NC_045512.2) using Minimap v2.12-r847 [34]. The BAM files were sorted by SAMtools sort [35]. The final sorted BAM files were used to call the genetic variants in variant call format (VCF) by SAMtools mpileup and BCFtools [35]. The final call set of the 30,983 genomes, was annotated and their impact was predicted using SnpEff v 4.3t [36]. For that, the SnpEff databases were first built locally using annotations of the reference sequence Wuhan-Hu-1/2019 obtained in the GFF format from NCBI database. Then, the SnpEff database was used to annotate SNPs and InDels with putative functional effects according to the categories defined in the SnpEff manual (http://snpeff.sourceforge.net/SnpEff_manual.html).

### D614G mutagenesis analysis

To investigate the possible impact of the most frequent D614G mutation, we conducted an *in-silico* mutagenesis analysis based on the CryoEM structure of the spike protein in its pre-fusion conformation (PDB id 6VSB). Modelling of the D614G mutation was done using UCSF Chimera [37]. Then, the mutant structure was relaxed by 1000 steps of steepest descent (SD) and 1000 steps of conjugate gradient (CG) energy minimizations keeping all atoms with more than 5A from G614 fixed. Comparative analysis of D614 (wild type) and G614 (mutant) interactions with their surrounding residues was done in PyMOL 2.3 (Schrodinger L.L.C).

### RBD mutations and Spike/ACE2 binding affinity

Modelling of RBD mutations was performed by UCSF chimera [37] using the 6M0J structure of the SARS-CoV-2 wild-type spike in complex with human ACE2 as a template. Mutant models were relaxed by 1000 steps of SD followed by 1000 steps of CG minimizations keeping all atoms far by more than 5A from the mutated residues fixed. Changes in the binding affinity of the spike/ACE2 complex for each spike mutant were estimated by the MM-GBSA method using the HawkDock server [38].

### Clustering and ivergence analysis

To evaluate the clustering with jaccrd distance between the genomes of each country based on their mutational frequencies, first the genomes were grouped in their countries of origin, then the mutation frequency was normalized individually (for each country), the matrix distance between countries was then measured using Jaccard Distance.

On the other hand, to calculate the intra-specific divergence of SARS-CoV-2, we used the Wuhan-Hu-1/2019 genome as a reference sequence, and the other 30,983 genomes were also sorted by country of origin.. The divergence was first calculated by estimating the similarities of the genomes with the reference sequence by grouping genomes from the same country using CD-Hit. [39]. All SARS-CoV-2 genomes used in this study were included except those from Ghana which were excluded from this analysis due to the high number of Ns. The percentage of similarity was then recovered to 100%. Then the percentage of divergence for each country was calculated using the following formula:

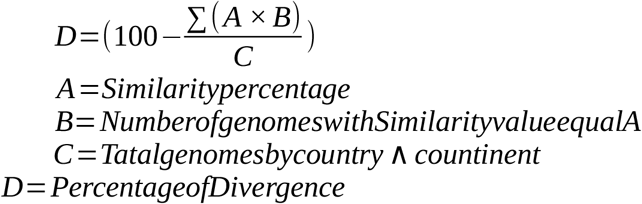

### Phylogenetic and spatio-dynamic analysis

We generated a phylogenetic and divergence tree, as well as a genomic epidemiology map based on the 30,983 genomes of SARS-CoV-2 using NextStrain tools (https://nextstrain.org) [40]. The tree was constructed in IQ-TREE v1.5.5 [41] using the maximum likelihood method under the GTR model. The rate of evolution and the Time to the Most Recent Common Ancestor (TMRCA) were estimated using ML dating in the Tree Time package [42].

## Conflict of interest

The authors declare that they have no competing interests.

## Acknowledgments

We sincerely thank the authors and laboratories around the world who have sequenced and shared the full genome data for SARS-CoV-2 in the GISAID database. All data authors can be contacted directly via www.gisaid.org.

This work was carried out under National Funding from the Moroccan Ministry of Higher Education and Scientific Research (Covid-19 Program) to AI. This work was also supported by a grant to AI from Institute of Cancer Research and the PPR-1 program to AI.

## Notes

### Competing Interest Statement

The authors have declared no competing interest.

